# Correlation of Infinium HumanMethylation450K and MethylationEPIC BeadChip arrays in cartilage

**DOI:** 10.1101/733204

**Authors:** Kathleen Cheung, Marjolein J. Burgers, David A. Young, Simon Cockell, Louise N. Reynard

## Abstract

**Background:** DNA methylation of CpG sites is commonly measured using Illumina Infinium BeadChip platforms. The Infinium MethylationEPIC array has replaced the Infinium Methylation450K array. The two arrays use the same technology, with the EPIC array assaying 865859 CpG sites, almost double the number of sites present on the 450K array. In this study, we compare DNA methylation values of shared CpGs of the same human cartilage samples assayed using both platforms.

**Methods:** DNA methylation was measured in 21 human cartilage samples using the Illumina Infinium Methylation450K BeadChip and the Infinium methylationEPIC array. Additional matched 450K and EPIC data in whole tumour and whole blood were downloaded from GEO GSE92580 and GSE86833 respectively. Data were processed using the Bioconductor package Minfi. Additionally, DNA methylation of six CpG sites was validated for the same 21 cartilage samples by use of pyrosequencing.

**Results:** In cartilage samples, overall sample correlations between methylation values generated by the two arrays were high (Pearson correlation coefficient r > 0.96). However, 50.5% of CpG sites showed poor correlation (r < 0.2) between arrays. Sites with limited variance and with either very high or very low methylation levels in cartilage exhibited lower correlation values, corroborating prior studies in whole blood. Bisulfite pyrosequencing did not highlight one array as generating more accurate methylation values that the other. For a specific CpG site, the array methylation correlation coefficient differed between cartilage, tumour and whole blood, reflecting the difference in methylation variance between cell types. These patterns can be observed across different tissues with different CpG site variances. When performing differential methylation analysis, the mean probe correlation co-efficient increased with increasing Δβ threshold used.

**Conclusion:** CpG sites with low variability within a tissue showed poor reproducibility between arrays. However, variance and thus reproducibility differs across different tissue types. Therefore, researchers should be cautious when analysing methylation of CpG sites that show low methylation variance within the cell type of interest, regardless of platform or method used to assay methylation.

## Introduction

Illumina methylation BeadChip arrays are a popular method for assaying methylation of CpG sites across the human genome. In 2016, the Illumina MethylationEPIC (EPIC) array replaced the Methylation450K (450K) array, which has since been discontinued. The EPIC array uses the same Infinium probe technology and share 452,453 common probes but assays almost double the number of CpG sites compared to the 450K array. The 450K and EPIC arrays utilise Infinium type I and type II probes. Type I probes consists of two probes, each binding to either methylated or unmethylated sites whereas type II probes use one probe to bind to the target CpG. Type I probes are reported to be more effective at assaying completely un-methylated or completely methylated probes [1]. The combination of type I and type II probes ensures that the whole range of methylation value can be assayed genome wide. The same CpG is assayed by the same probe type on both arrays; sites do not change probe chemistries between arrays.; a comprehensive description of probe types is available [2]. As ongoing studies and meta-analyses may use data from both array platforms, previous studies have assessed the concordance of the two by correlating methylation levels of the same samples assayed on both arrays. Studies comparing the two arrays have been reported in tumour, whole blood and placental samples [3–6]. Whilst high overall per sample correlations (Pearson’s *r* > 0.9) were reported, two of the blood studies found that many individual sites in whole blood showed poor correlations (*r* < 0.2) [2,3]. The studies in whole blood suggested that CpG sites with low variance and that are either completely methylated or un-methylated tend to exhibit poor reproducibility between 450K and EPIC arrays. Furthermore, several studies have highlighted sites with low reproducibility in blood and placental tissue that they have recommended be removed from the analysis when performing meta-analysis of 450K and EPIC datasets [3–5].

In this study, we investigate the correlation of the 450K and EPIC platforms in human cartilage, a tissue which comprises a single cell type. As variance of DNA methylation in single CpG sites has been found to affect correlation across 450K and EPIC arrays, we hypothesise that tissue types with different inherent DNA methylation variances do not have the same reproducibility across the two arrays. Tumour tissue is heterogeneous and whole blood consists of many cell types, factors which affect the variance of DNA methylation of CpG sites in a sample. We assayed 22 cartilage samples on both BeadChip arrays and examined per sample and per CpG correlations. We also downloaded publicly available 450K and EPIC data in whole tumour (GEO GSE92580) and whole blood (GSE86833) to compare the variance and correlation across arrays with cartilage samples. We sought to investigate the accuracy of array generated methylation values for six CpG sites using the gold standard of bisulfite pyrosequencing in the 21 samples common to both arrays.

## Results

### Overall sample correlation between 450K and EPIC arrays

DNA methylation in human cartilage samples, nine from patients undergoing total joint replacement with osteoarthritis (OA) of the knee and 12 with neck of femur fracture (NOF) were assessed using the 450K and EPIC arrays. The correlation between the same samples ran on both arrays were examined, with probes common to both arrays included in the analysis. In total, 450K and EPIC share 452,453 probes. Quality control measures and probe filters including removal of sex chromosome and cross reacting probes were applied to ensure only probes which could be measured with confidence were used for further analysis [See methods section]. After quality control of probes, 370,669 remained. Additionally, all cartilage samples had > 95% probes with a detection *p* value < 0.01 on both 450K and EPIC arrays. Both arrays were normalised using the functional normalisation method [7], which uses control probes to account for unwanted variation in the data.

To investigate the correlation between the arrays, the Pearson correlation coefficient was calculated per sample for the 370,699 probes common to both arrays and passing quality filters. Corroborating previous studies [3,4,6,8], we found that the per sample correlation of methylation β-value was high. A 450K vs EPIC scatter plot of methylation β-value for one cartilage sample illustrates that the overall correlation was high (Pearson’s *r* 0.99) although many CpGs show greater than 5% difference between arrays (Fig. 1A). The overall per sample correlation between the same sample assayed on the EPIC array twice was also high at *r* = 0.99 (Fig. 1B). However, similar to Logue et al [3], we also observed that pairwise correlations between any two samples from different individuals and arrays were high. A correlation heatmap showed that samples were more likely to be similar to other samples assayed on the same array, rather than to itself run on the other array (Fig. 1C). Separation between arrays is also seen and is likely a batch effect; nevertheless, we still observe high Pearson’s *r* across all samples which implies that a high overall correlation may not necessarily reflect biological similarity.

**Figure 1.**
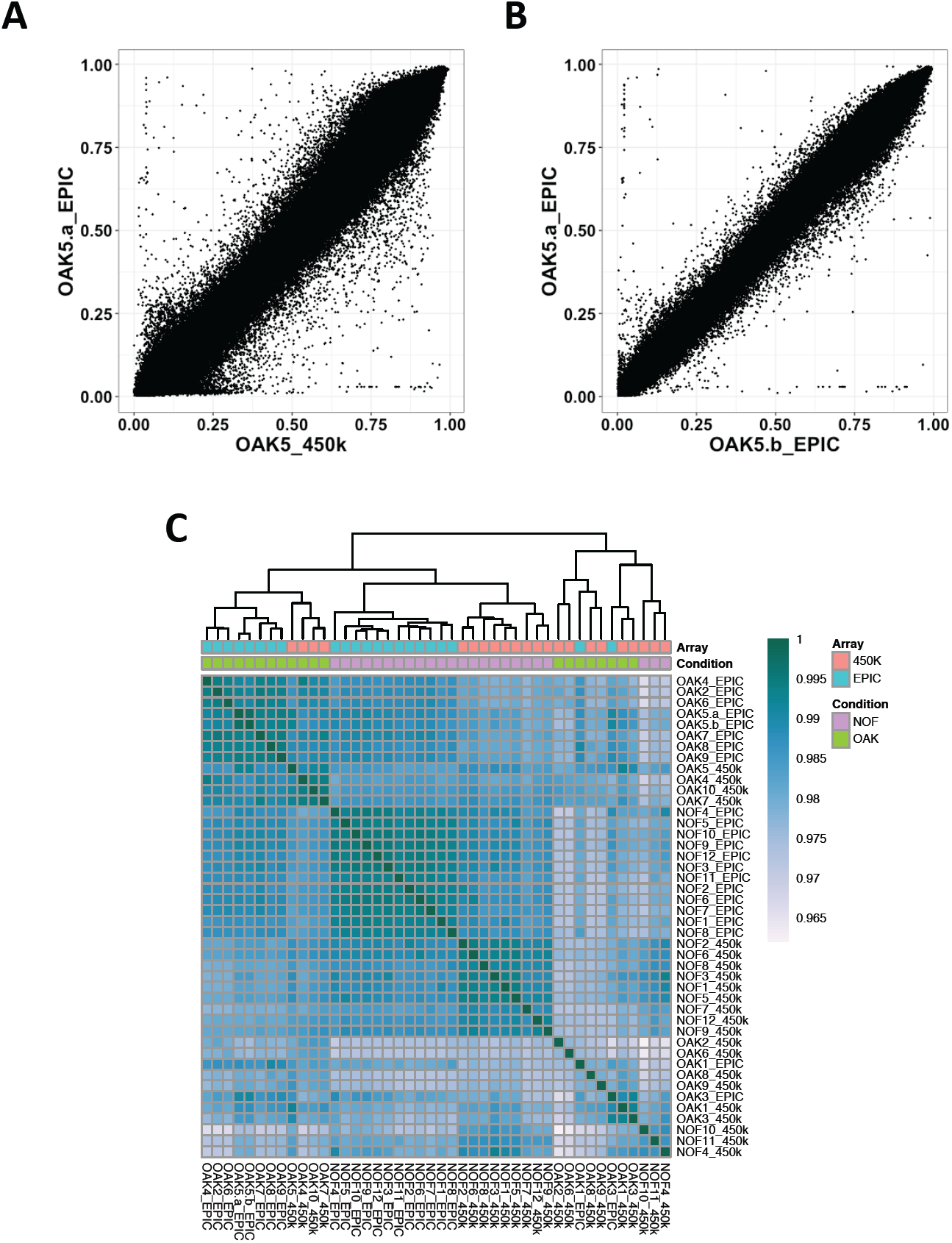
Correlation of normalized methylation β-values in 450K vs EPIC arrays. (A) Correlation of β values between the OAK5 cartilage sample assayed on the 450K array and the EPIC array (Pearson’s r = 0.99). (B) β value correlation of the same sample (OAK5) assayed twice on the EPIC array (Pearson’s r = 0.99). (C) β value correlation heatmap of all 21 cartilage samples assayed on both the 450K and EPIC array.

### Variance and correlation of CpG sites by probe type

We next examined the correlation between 450K and EPIC generated β values for individual CpG sites common to the 450K and EPIC arrays. Similar to prior studies, we found that over 50% of sites showed poor correlation (Pearson’s *r* <0.2), with a median per CpG correlation of 0.195 (mean 0.218) (Supplemental Table 1A). Only 3.56% of probes showed high (≥0.8) correlation values. We also observed that CpG site methylation variance was generally low across our cartilage samples (Fig.2A). Poor correlation CpG sites (*r* < 0.2) had a significantly lower β value variance than high correlation (*r* > 0.8) sites (median variance of 0.0018 and 0.0064 respectively, p<2.2×10^-16^, Fligner-Killeen non-parametric variance test) This confirms that the pattern of low variance sites and poor correlation observed in whole blood [3,4] and placenta [5] is also seen in cartilage.

We sought to determine whether there were differences in CpG variance and correlation assayed by probe types in cartilage, as has been reported in blood [3]. Solomon et al [4] found that type II probes had a larger proportion of high correlation probes in comparison to type I. The 450K array consisted of 28% Infinium type I probes whereas the EPIC array consists of 16% type I probes. We observed that CpG sites assayed using Infinium type I probes had significantly lower variance compared to sites measured with type II probes (Fig. 2B; p<2.2×10^-16^, Fligner-Kileen test). Furthermore, there was a trend for type I probes =to show lower CpG correlations compared to type II (Fig. 2C).

**Figure 2.**
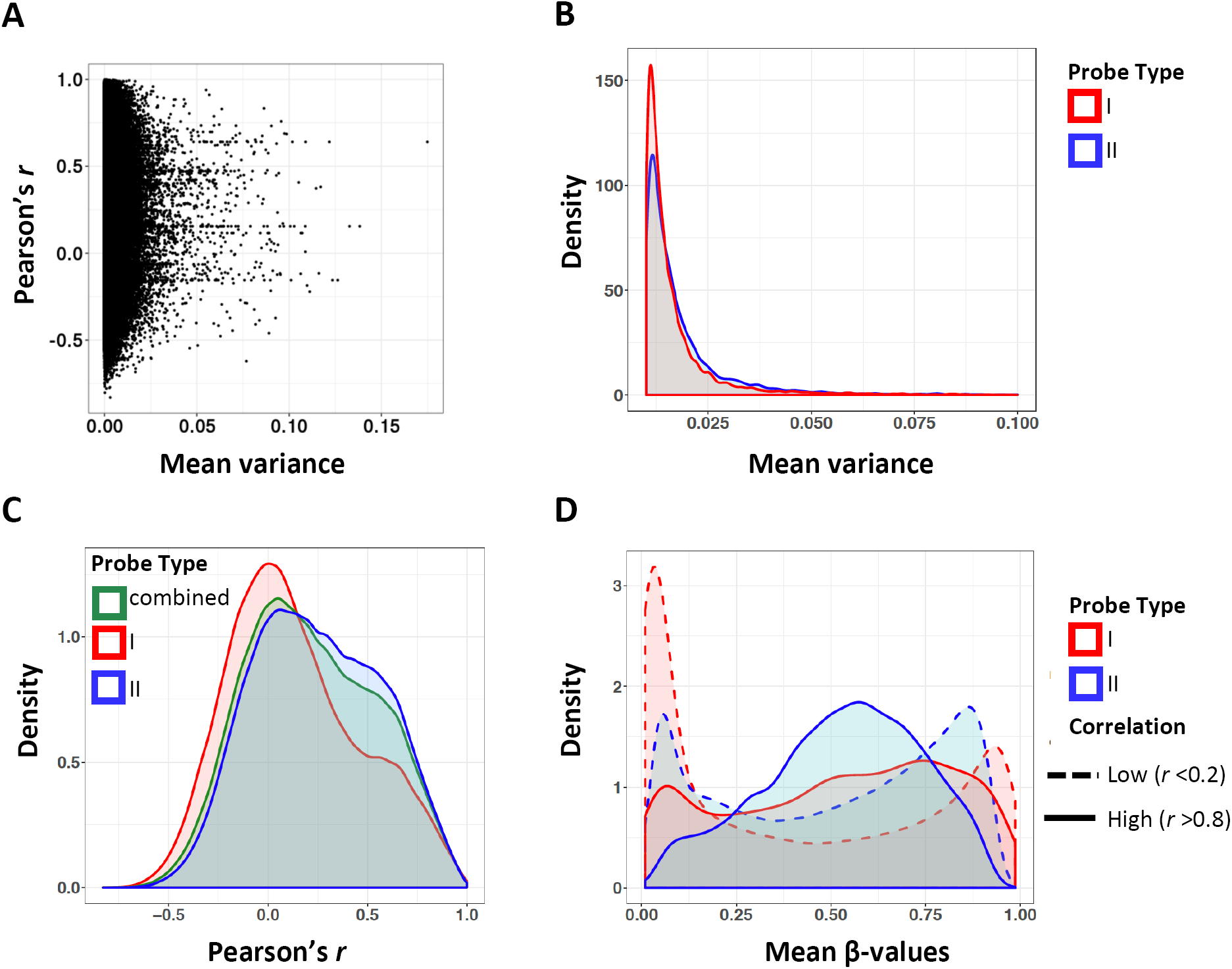
Correlation and variance of methylation β-values by probe type. (A) Methylation (β value) variance plotted against Pearson correlation coefficient across common CpG sites in 450K and EPIC arrays. Variance is the mean of the 450K and EPIC variances across the 21 samples. (B) Density of β value variance by probe type. Type I probes are shown in red and type II probes are in blue. (C) Density of Pearson correlation coefficient *r* by probe type. Type I probes are red, type II probes are in blue and the combined Pearson coefficient of both probe types are in green. (D) Mean methylation β-values by probe type and correlation group. CpG sites with low correlation (r < 0.2) are denoted by a dashed line whereas high correlation (r > 0.8) are marked with solid lines.

It has previous been observed that CpG sites with methylation levels at or close to extremes (either 0% or 100% methylation) were more prone to low variance and poor reproducibility [2,3]. We determined the mean methylation β-values per CpG in four groups: probe type I and high correlation, probe type I and low correlation, probe type II and high correlation, probe type II and low correlation (Fig. 2D). In the low correlation groups, regardless of probe chemistry, methylation β levels were indeed skewed towards 0% or 100%. This skew is more pronounced in type I probes, although the same trend is also seen in type II probes.

### Differences in variance and correlation between tissue types

To examine if there are differences in DNA methylation variance and array β value correlations between tissue types, we compared our cartilage correlations with those of previously published 450K vs EPIC datasets. The raw data from matched 450K and EPIC samples collected from fresh-frozen paediatric brain tumours and whole blood from Guthrie cards were accessed from GEO GSE92580 and GSE86833 respectively. Data were filtered and processed in the same way as our cartilage datasets.

The Guthrie card whole blood dataset showed similar median (*r* = 0.285) and mean (*r* = 0.221) individual CpG correlations to cartilage (Supplemental Table 1A) and other whole blood comparisons [3,4], with only 16.68% of probes having Pearson correlations above 0.8 (Fig. 3A, Supplemental Table 1A). Interestingly, in contrast to whole blood and cartilage, the majority (58. 60%) of individual CpG sites within the tumour dataset exhibited high correlation (≥0.8) between the two arrays, with a median Pearson’s *r* of 0.888 and a mean of 0.695 (Fig. 3A, B-, Supplemental Table 1A). We further observed that variance of individual CpG sites in tumour were higher than in whole blood and cartilage (Fig. 3C). Together, our data supports the hypothesis that low methylation variance sites tend to have poorly reproducible β values between the two arrays, a pattern we observe across both probe types and across different tissue types.

**Figure 3.**
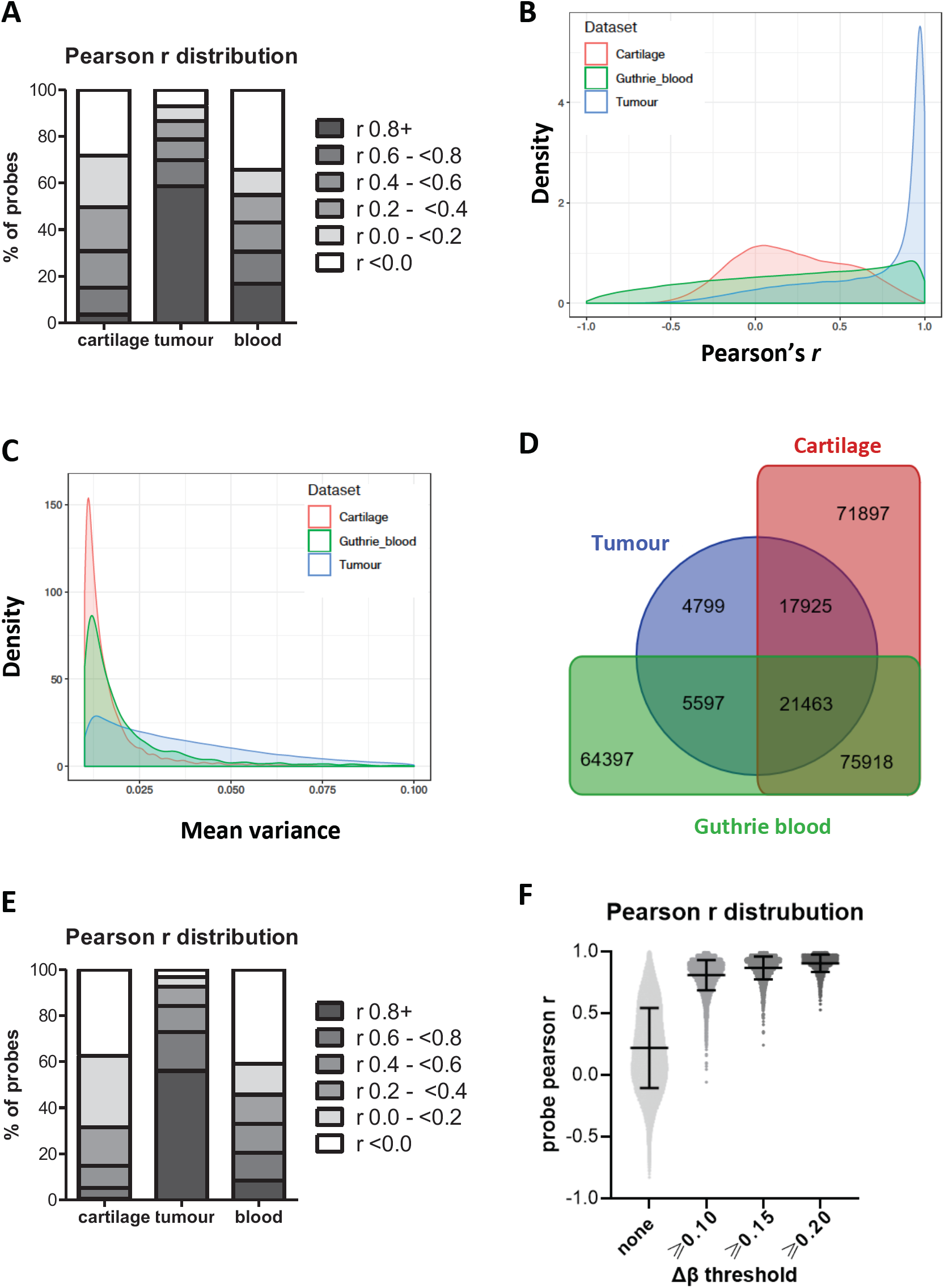
Pearson correlation and β value variance of CpG sites across 450K and EPIC arrays in three tissue types. Common samples assayed on both 450K and EPIC arrays were investigated in whole blood (GEO GSE86833) and pediatric brain tumour (GEO GSE92580). (A) Percentage of probes grouped by Pearson’s *r* values. (B) Density plot of Pearson’s *r* of CpG sites in cartilage (red), whole blood (green) and whole tumour (blue). (C) Density plot of mean variance (450K and EPIC) in cartilage, whole blood and whole tumour. (D) Venn diagram of low correlation (*r* < 0.2) sites in cartilage, tumour and whole blood. (E) The grouped distribution of Pearson’s *r* values in cartilage, tumour and whole blood of low correlating probes (r ≤0.2) identified in 108 placental samples by Fernandez-Jimenez and colleagues (ref). (F) Pearson *r* distribution of differentially methylated CpGs between OA knee and NOF cartilage samples plotted against Δ β thresholds. The cartilage *r* distribution for all probes is shown on the left for comparison.

Given the difference in Pearson *r* distribution and mean β variance between cartilage, tumour and whole blood, we investigated the degree of overlap of high (≥0.8) and poorly correlating probes (r <0.2) in the three tissues (Fig. 3D, Supplemental Table 1B). For a specific CpG site, DNA methylation variance is different between the three tissue types analysed, such that CpG sites with low 450K-EPIC correlations are different between the tissue types (Supplemental Table 1B). Of 261,995 CpG sites that show low (r<0.2) β value correlations between the two arrays in at least one tissue, only 8.19% were common between them (Fig. 3D). As low correlation sites are mostly distinct between these tissue types, researchers should not extrapolate data on 450K-EPIC probe correlations from one tissue to inform another study using different tissue samples. This is further confirmed by examining the Pearson r distribution in cartilage, tumour and blood of 2668 probes recently reported to show poor correlations (r ≤0.2) between arrays in 108 placental samples [5]. Although more than half of these low reproducibility placental probes also poorly correlate across the 450K and EPIC arrays in cartilage and blood samples, the majority (56.07%) show Pearson correlations of ≥0.8 in tumour (Fig. 3E, Supplemental Table 1C).

### Comparison with pyrosequencing

It has suggested that the methylation values generated by the EPIC array are more accurate than those generated by 450K array as the former contains more type II probes [4]. To investigate this, we measured DNA methylation of six CpG sites that are present on both arrays using bisulfite pyrosequencing. Pyrosequencing is a technique that uses sequencing-by-synthesis to determine nucleotide polymorphisms in DNA and can be used to quantify DNA methylation after bisulfite conversion of DNA [7]. The six CpGs were selected to reflect the range of β value correlations observed in cartilage; two with negative correlations and very different β values between the two arrays (Supplemental Fig. S1), three with intermediate *r* values (*r* 0.430 to 0.711) and one with high correlation (cg13782176, *r* = 0.976). After confirming the accuracy of pyrosequencing DNA methylation values using DNA of known methylation levels, we measured the same 21 cartilage DNA samples that we had matched 450K and EPIC data for (Supplemental Fig. S1 and S2). Our limited pyrosequencing validation was inconclusive for deducing which array platform generated more accurate methylation values; for the two CpGs that were non concordant between 450K and EPIC arrays, the pyrosequencing methylation values for one (cg24555089; Fig. S1A) was more similar to the 450K values and the other (cg20806296; Fig. S1B) was more similar to the EPIC values. CpG sites with similar methylation β-values between 450K and EPIC also yielded similar methylation values measured by pyrosequencing (Fig. S1C). Prior studies examining the concordance between the 450K array and pyrosequencing have found that, whilst showing a good overall correlation, measurements of some CpG sites do not replicate between the two methods [9].

For epigenome wide association studies (EWASs), the ability to combined 450K and EPIC generated DNA datasets to perform meta-analyses is of great interest. The majority of EWASs use both a Δβ threshold and p value threshold for identifying differentially methylated sites (DMSs), although the Δβ thresholds vary widely between studies. We performed differential methylation analysis of 12 NOF cartilage samples and nine OA knee samples in order to examine if changes in Δβ threshold altered the probe Pearson r distribution and thus reproducibility between the 450K and EPIC arrays (Fig. 3F, Supplemental Table 1D). For DMSs identified by both arrays, the mean and median Pearson *r* increases with increasing Δβ threshold (Fig 3.F), presumably due to selection of probes increasingly variable methylation levels. Furthermore, the percentage of highly correlating probes (*r* ≥0.8) rising from 61.90% to 90.78% when the Δβ threshold is raised from 0.1 to 0.2. Thus, when performing a meta-analysis of data generated by the two arrays, the reproducibility between the datasets will increase with increasing Δβ threshold used for identifying DMSs.

## Discussion

We have shown that whilst overall correlation is high between the same cartilage samples assayed on 450K and EPIC arrays, some individual CpG sites show poor reproducibility. We found that type I probes are more prone to low correlation than type II. Type I probes are more effective than type II probes at measuring the extremes of methylation (0% or 100%) but it is at these extremes that CpG sites show poor reproducibility. Studies in whole blood also reported poor individual CpG site correlations [3,4]. These blood studies used different normalisation procedures suggesting that this artefact is present regardless of pre-processing pipeline. Furthermore, this study encompasses fewer samples compared to previously published comparisons and the conclusions remain the same.

Comparisons across tissues reveals that this trend of poor reproducibility in low variance sites is likely due to the inherent variability of DNA methylation in tissue or cell types rather than analysis method. Cartilage tissue has a lower DNA methylation variance in comparison to whole tumour and whole blood. This could be in part due to the composition of cell types within these tissues; whole blood is comprised of numerous cell types and tumour tissue consists of normal and abnormal cells, factors which contribute to heterogeneity in DNA methylation [10–12]. A study comparing 450K and EPIC in placental tissue, a tissue comprised of multiple cell types, found that per CpG correlations were high [5]. CpG sites with poor reproducibility vary between tissue types and therefore the specification of a common list of CpGs to exclude would not be applicable. Researchers studying DNA methylation in tissues or cell types exhibiting low variance or extreme methylation values should be aware that some CpG sites may not be reproducible.

The 450K array is no longer available; researchers wishing to assay methylation using BeadChip technology only have the option of the EPIC array. Although the 450K array has been discontinued by Illumina, the array may still have relevance in ongoing studies and meta-analyses. The comparisons between the 450K and EPIC has revealed an interesting technical artefact in measuring the methylation of low variance sites with either 0% or 100% methylation levels. However, it is not the first time that poorly correlated CpG sites have been described. Comparisons between the 450K and its predecessor, the Infinium HumanMethylation27 BeadChip showed strong overall correlations but weak correlation of some CpG sites [1,13]. Past studies examining the similarity of measurements in 450K and pyrosequencing in cell lines and brain tissue [9,14] have also observed that some CpG sites do not reproduce. Moreover, comparisons of 450K to whole genome bisulfite sequencing showed the same conclusions [1,15,16] as did reduced representation bisulfite sequencing [17,18]. Multiple published manuscripts have described the correlation of CpG sites measured using different methods. These studies were performed in various cell lines and tissue types. All report a high overall correlation between DNA methylation of the same samples measured using different methods. Not all of these studies commented on individual CpG site correlation, however, manual inspection of the correlation scatter plots reveal that a subset of CpGs show large differences in methylation values between the platforms being compared [1,15–18]. Altogether, these studies demonstrate that some CpGs reproduce poorly, regardless of sample type or method used.

It is not often feasible for researchers to validate CpGs genome-wide using a different method and correlate between methods in order to find sites with poor reproducibility. However, we and others have shown that CpG sites with poor correlation also tend to be those with low variance and very high or very low methylation. Therefore, it may be beneficial to apply a filter based on variance and/or percentage methylation especially where the methylation level of individual CpGs is important for downstream analysis or if DNA methylation in the tissue type being studied is relatively homogeneous. For example, studies investigating the effect of single CpG sites such as methylation quantitative trait loci (mQTL) or epigenome-wide association studies (EWAS) may wish to apply additional filters. Whilst researchers may wish to validate potentially important CpG sites using a different assay, some sites may not reliably reproduce regardless of method. Additionally, it can be difficult to reliably deduce which method shows the correct methylation level.

To conclude, we have corroborated past studies comparing the reproducibility between the 450K and EPIC platforms. We found that whilst per sample correlation was high, correlation between individual CpGs tended to be low in cartilage. We also report that CpG sites with poor reproducibility differ between tumour, whole blood and cartilage and this could be due to differences in DNA methylation variance between tissues.

## Methods

### Collection of cartilage samples and DNA methylation analysis

Cartilage samples were obtained from patients with knee OA (n=9) or neck of femur fracture (NOF, n=12) undergoing total joint replacement surgery at the Newcastle upon Tyne NHS Foundation Trust hospitals. The Newcastle and North Tyneside Research Ethics Committee granted ethical approval for the collection of the samples (REC reference number 14/NE/1212), with each donor providing informed consent. Cartilage was ground as previously described (Raine et al 2012 PMID 22586168) and DNA extracted using the E.Z.N.A Tissue DNA kit (omega BIO-TEK). 1ug of DNA was bisuphite converted using the EZ DNA methylation kit (Zymo Research) and quantified using the 450K and EPIC arrays by Edinburgh Clinical Research Facility.

### Analysis of 450K and EPIC data

Data were processed using the Bioconductor package *minfi* [19]. Samples were normalised using the functional normalisation method implemented in the *minfi* package. All samples had > 95% probes passing a detection p value of 0.01. Probes with a detection *p* value < 0.01 were removed from the analysis. Probes within sex chromosomes were removed from the analysis, as were probes previously reported to be cross-reacting [20] and probes containing SNPs with a minimum allele frequency > 0.05 in European populations. Only probes common to the 450K and EPIC array were selected, this left 372278 probes (96514 Type I and 275764 Type II) for analysis. Age and sex covariates of cartilage samples were corrected for using the removeBatchEffect function in *limma* [21]. Correction was performed using methylation M-values before conversion to methylation β-values. Pearson correlation coefficient for was calculated per sample and per CpG site across both 450K and EPIC arrays. Datasets GEO GSE92580 and GSE86833 were analysed using exactly the same method. Correlation and density plots were generated using the ggplot2 package in R. An online webtool was used to create the Venn diagram (http://bioinformatics.psb.ugent.be/webtools/Venn/).

### Targeted pyrosequencing

Genomic DNA (750 ng) of the same 21 cartilage samples was bisulphate converted as above and eluted in 40 µl elution buffer. Pyrosequencing assays for the six selected CpGs were designed using PyroMark assay design software 2.0 (Qiagen), the primers used are listed in Supplementary Table S2. Pyrosequencing was performed using PyroMark Q24 (Qiagen). A calibration curve was made to confirm accuracy of the pyrosequencer. For each CpG, two gBlocks (IDT) of the 300bp region surrounding the CpG site of interest were designed to the bisuphite sequence with either the CpG site methylated (C) or unmethylated (T), then the pyrosequencing calibration curve was performed using input DNA of 0-100% methylation with 10% increments.

## Declarations

### Authors’ contributions

LNR, DAY, SC and KC designed the study. LNR and MJB collected the data. KC performed the data analysis. All authors read and approved the manuscript.

**Supplemental Figure 1.**
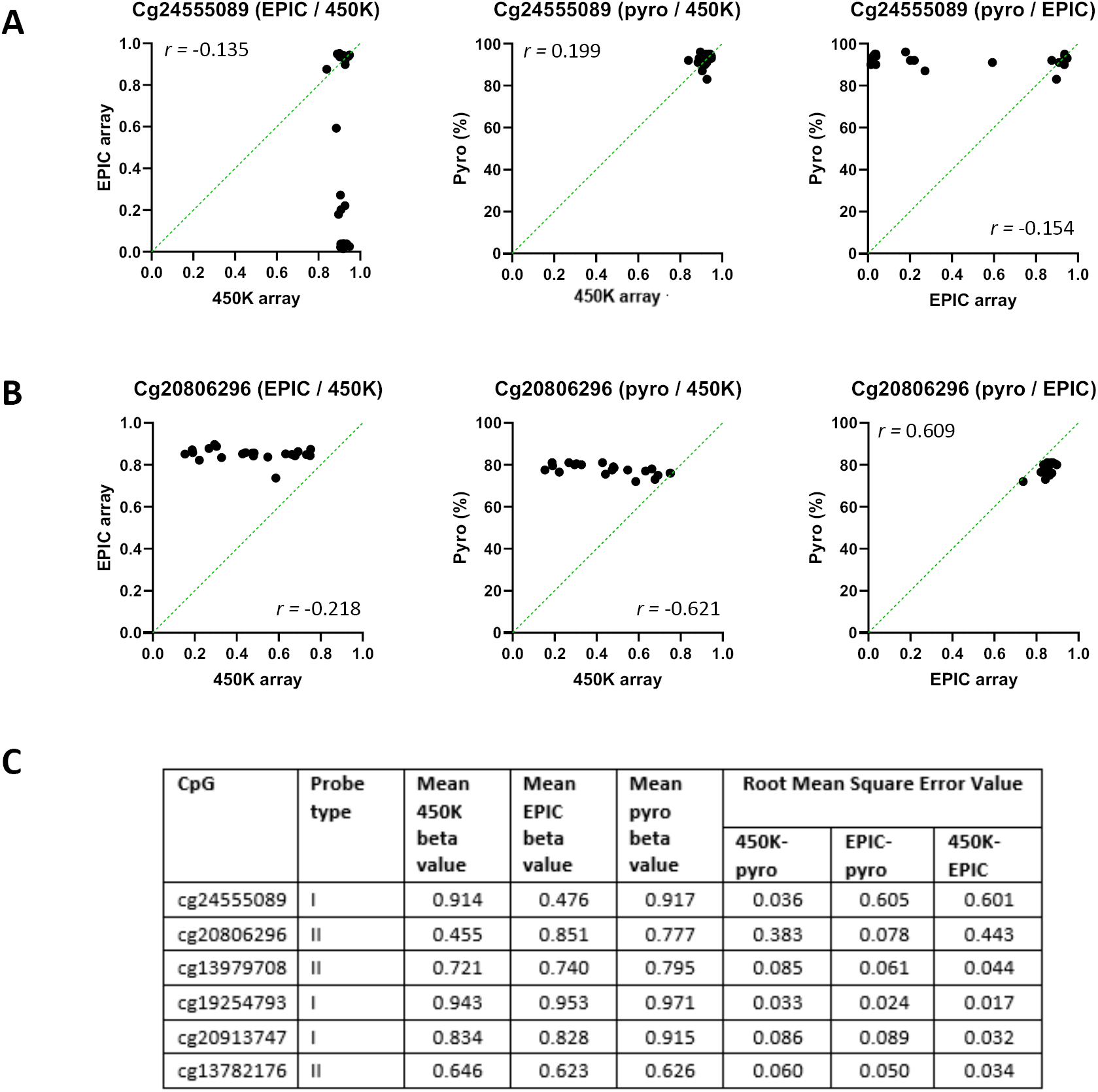
Comparison of methylation levels in 21 cartilage samples measured by 450K array, EPIC array, and pyrosequencing for six CpG sites. (A) and (B) Methylation levels (A) cg24555089 and (B) cg20806296, both of which have very different β values for the same 21 samples between the 450K and EPIC arrays. Left graph, 450K vs EPIC β value; middle graph, 450K vs pyrosequencing % methylation; right graph, EPIC vs pyrosequencing % methylation. The dotted green line represents when the methylation levels detected by the two techniques are the same. The accuracy of each pyrosequencing assay was validated using a calibration curve of known methylation values of 0-100%. (C) Table of the mean methylation levels measured by the two arrays and pyrosequencing for all six CpGs assayed. β value is given for the two arrays, and the percentage methylation measured by pyrosequencing has been converted to a β value. The Root Mean Square Error Value is given for the pairwise comparison between two methods; the smaller the value, the more similar the β values quantified by the two methods are.

**Supplemental Figure 2.**
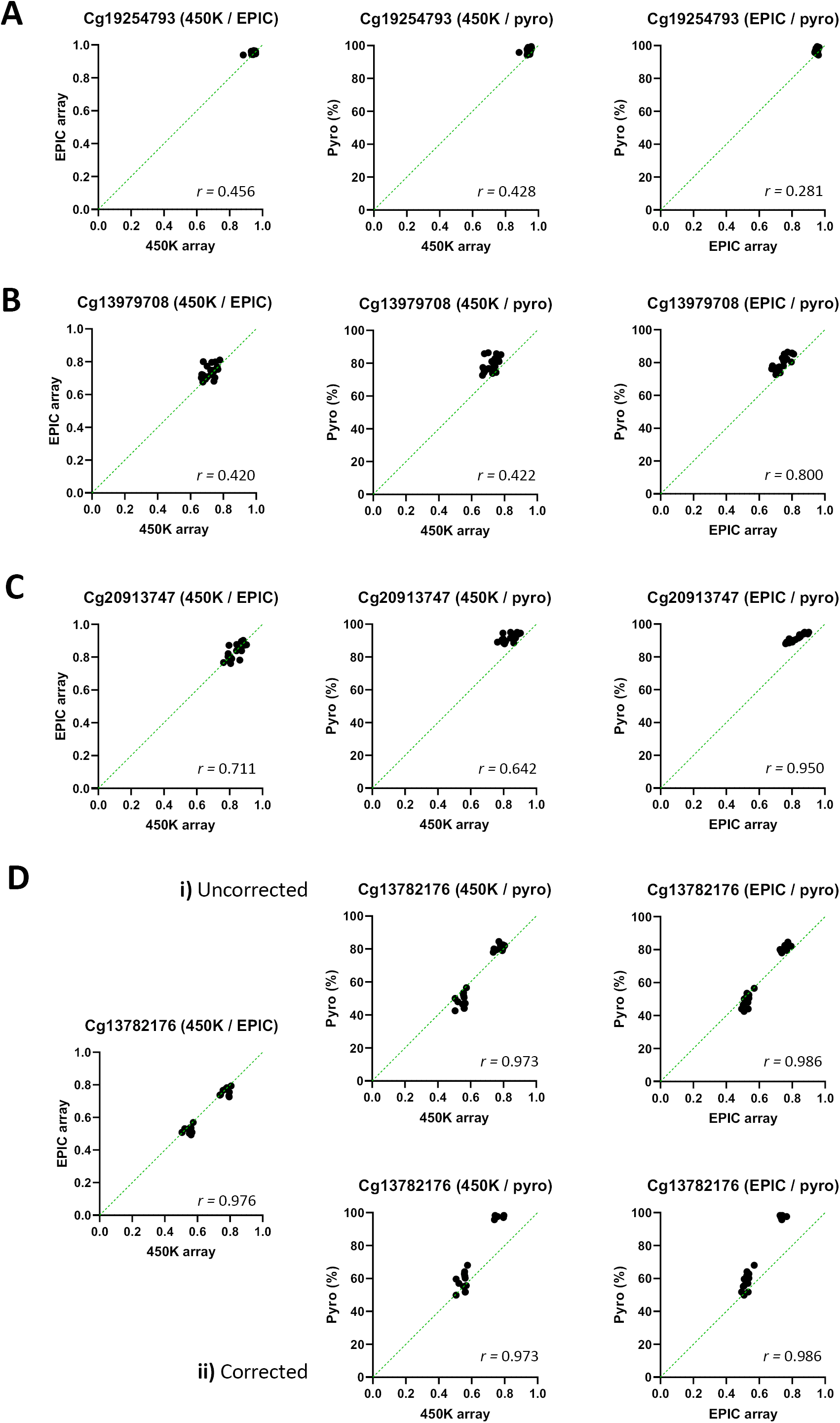
Comparison of methylation levels in 21 cartilage samples measured by 450K array, EPIC array, and pyrosequencing for (A) cg19254793, (B) cg1397908, (C) cg20913747 and (D) cg13782176. Left graph, 450K vs EPIC β value; middle graph, 450K vs pyrosequencing % methylation; right graph, EPIC vs pyrosequencing % methylation. The dotted green line represents when the methylation levels detected by the two techniques are the same. The accuracy of each pyrosequencing assay was validated using a calibration curve of known methylation values of 0-100%. Only the cg13782176 calibration curve deviated from the expected value, and so both the (i) uncorrected and (ii) corrected pyrosequencing methylation levels for cg13782176 are shown in (D).

**Supplemental Table 1.**
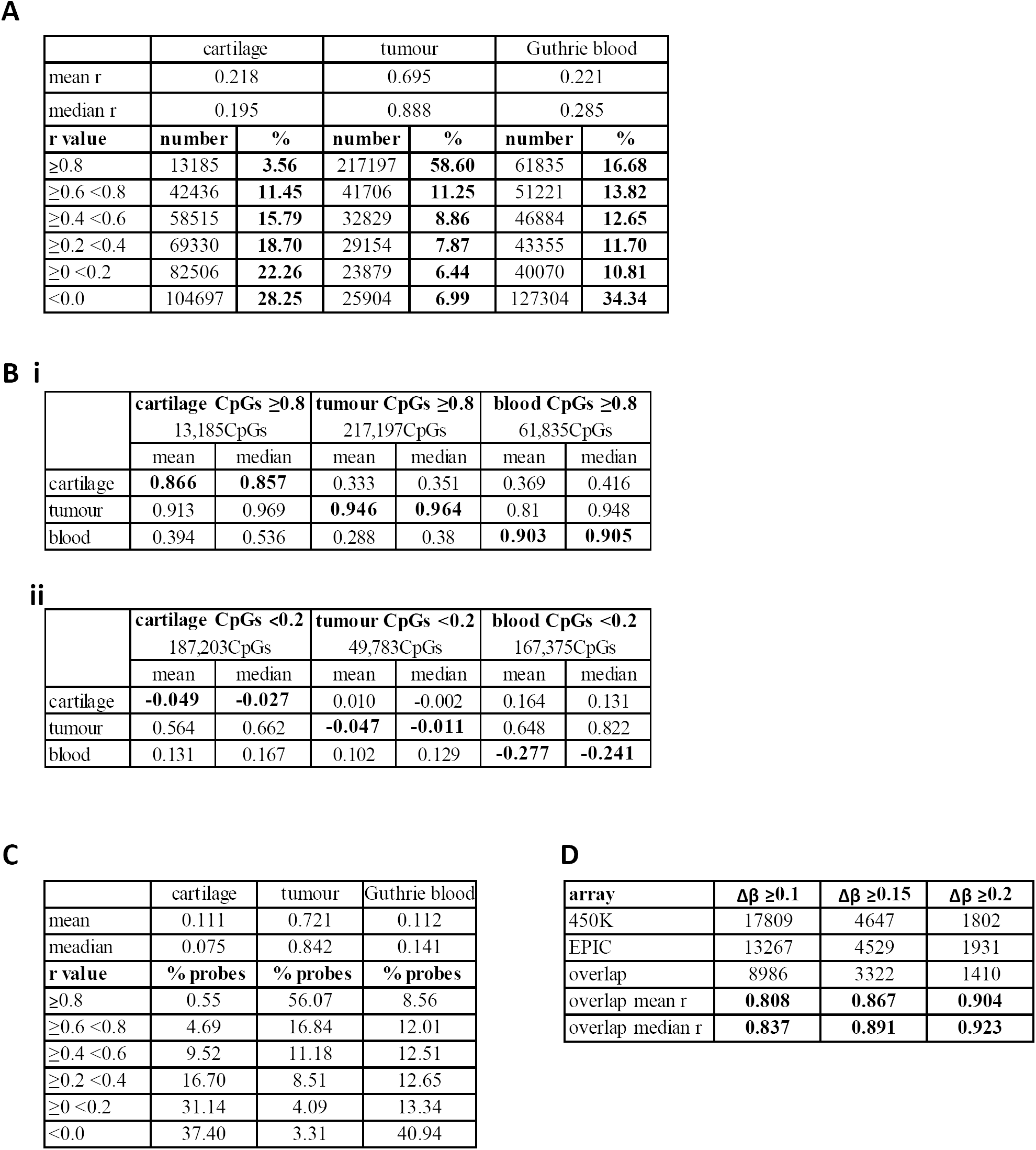
Correlation of CpG sites across 450K and EPIC arrays in three tissue types. Common samples assayed on both 450K and EPIC arrays were investigated in whole blood (GEO GSE86833) and pediatric brain tumour (GEO GSE92580). (A) Percentage of total numbers of CpG probes grouped by Pearson’s *r*. (B) Mean and median Pearson *r* values per tissue for (i) CpGs with *r* >0.8 and (ii) *r* <0.2 in at least one tissue. (C) The grouped distribution of Pearson’s *r* values in cartilage, tumour and whole blood of low correlating probes (r ≤0.2) identified in 108 placental samples by Fernandez-Jimenez and colleagues (ref). (D) Number of differentially methylated CpGs identified between NOF and OA knee cartilage using different Δβ thresholds. For overlapping CpGs identified by both 450K and EPIC arrays, the mean and median Pearson r is shown.

|  |  |  |  |  |  |  |
| --- | --- | --- | --- | --- | --- | --- |
|  | cartilage |  | tumour |  | Guthrie blood |  |
| mean r | 0.218 |  | 0.695 |  | 0.221 |  |
| median r | 0.195 |  | 0.888 |  | 0.285 |  |
| r value | number | % | number | % | number | % |
| ≥0.8 | 13185 | 3.56 | 217197 | 58.60 | 61835 | 16.68 |
| ≥0.6 <0.8 | 42436 | 11.45 | 41706 | 11.25 | 51221 | 13.82 |
| ≥0.4 <0.6 | 58515 | 15.79 | 32829 | 8.86 | 46884 | 12.65 |
| ≥0.2 <0.4 | 69330 | 18.70 | 29154 | 7.87 | 43355 | 11.70 |
| ≥0 <0.2 | 82506 | 22.26 | 23879 | 6.44 | 40070 | 10.81 |
| <0.0 | 104697 | 28.25 | 25904 | 6.99 | 127304 | 34.34 |
| B i |  |  |  |  |  |  |
|  | cartilage CpGs ≥0.8 |  | tumour CpGs ≥0.8 |  | blood CpGs ≥0.8 |  |
|  | 13,185CpGs |  | 217,197CpGs |  | 61,835CpGs |  |
|  | mean | median | mean | median | mean | median |
| cartilage | 0.866 | 0.857 | 0.333 | 0.351 | 0.369 | 0.416 |
| tumour | 0.913 | 0.969 | 0.946 | 0.964 | 0.81 | 0.948 |
| blood | 0.394 | 0.536 | 0.288 | 0.38 | 0.903 | 0.905 |
|  | cartilage CpGs <0.2 |  | tumour CpGs <0.2 |  | blood CpGs <0.2 |  |
|  | 187,203CpGs |  | 49,783CpGs |  | 167,375CpGs |  |
|  | mean | median | mean | median | mean | median |
| cartilage | -0.049 | -0.027 | 0.010 | -0.002 | 0.164 | 0.131 |
| tumour | 0.564 | 0.662 | -0.047 | -0.011 | 0.648 | 0.822 |
| blood | 0.131 | 0.167 | 0.102 | 0.129 | -0.277 | -0.241 |

|  |  |  |  |
| --- | --- | --- | --- |
|  | cartilage | tumour | Guthrie blood |
| mean | 0.111 | 0.721 | 0.112 |
| meadian | 0.075 | 0.842 | 0.141 |
| r value | % probes | % probes | % probes |
| ≥0.8 | 0.55 | 56.07 | 8.56 |
| ≥0.6 <0.8 | 4.69 | 16.84 | 12.01 |
| ≥0.4 <0.6 | 9.52 | 11.18 | 12.51 |
| ≥0.2 <0.4 | 16.70 | 8.51 | 12.65 |
| ≥0 <0.2 | 31.14 | 4.09 | 13.34 |
| <0.0 | 37.40 | 3.31 | 40.94 |

|  |  |  |  |
| --- | --- | --- | --- |
| array | Δβ ≥0.1 | Δβ ≥0.15 | Δβ ≥0.2 |
| 450K | 17809 | 4647 | 1802 |
| EPIC | 13267 | 4529 | 1931 |
| overlap | 8986 | 3322 | 1410 |
| overlap mean r | 0.808 | 0.867 | 0.904 |
| overlap median r | 0.837 | 0.891 | 0.923 |

---

**Supplemental Table 2.**
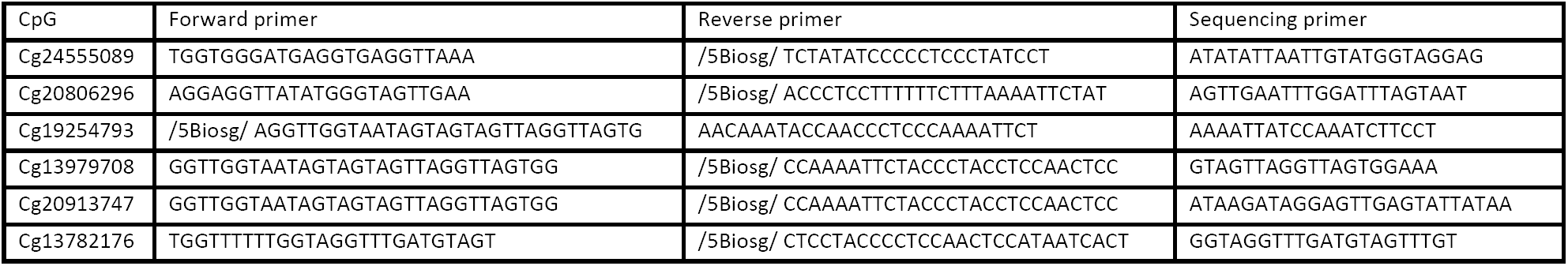
Pyrosequencing primerss.

